# Rethinking the Methods and Algorithms for Inner Speech Decoding - and Making them Reproducible

**DOI:** 10.1101/2022.03.22.485286

**Authors:** Foteini Simistira Liwicki, Vibha Gupta, Rajkumar Saini, Kanjar De, Marcus Liwicki

## Abstract

This study focuses on the automatic decoding of inner speech using noninvasive methods, such as electroencephalography (EEG)). While inner speech has been a research topic in philosophy and psychology for half a century, recent attempts have been made to decode nonvoiced spoken words by using various brain-computer interfaces. The main shortcomings of existing work are reproducibility and the availability of data and code. In this work, we investigate various methods (using Convolutional Neural Network (CNN), Gated Recurrent Unit (GRU), Long Short-Term Memory Networks (LSTM)) for the detection task of 5 vowels and 6 words on a publicly available EEG dataset. The main contributions of this work are (1) subject dependent vs. subject-independent approaches, (2) the effect of different preprocessing steps (Independent Component Analysis (ICA), down-sampling and filtering), and (3) word classification (where we achieve state-of-the-art performance on a publicly available dataset). Overall we achieve a performance accuracy of 35.20% and 29.21% when classifying 5 vowels and 6 words respectively in a publicly available dataset, using our tuned iSpeech-CNN architecture. All of our code and processed data are publicly available to ensure reproducibility. As such, this work contributes to a deeper understanding and reproducibility of experiments in the area of inner speech detection.

## 0. Introduction

Thought is strongly related to inner speech [1,2], through a voice being inside the brain that does not actual speak. Inner speech, although not audible, occurs when reading, writing, and even when idle (i.e., “mind-wandering” [3]). Moreover, inner speech follows the same pattern e.g., regional accents, as if the person is actually speaking aloud, for example [4]. This work focuses on inner speech decoding.

While inner speech has been a research topic in the philosophy of psychology since the second half of the 20th century [5], with results showing that the part of the brain responsible for the generation of inner speech is the frontal gyri, including Broca’s area, the supplementary motor area and the precentral gyrus, the automatic detection of inner speech has very recently become a popular research topic [6], [7]. However, a core challenge of this research is to go beyond closed vocabulary decoding of words and integrate other language domains (e.g., phonology and syntax) to reconstruct the entire speech stream.

In this work, we conduct extensive experiments using deep learning methods to decode 5 vowels and 6 words on a publicly available electroencephalography (EEG) dataset [8]. The backbone CNN architecture used in this work is based on the work of Cooney et al. [7].

The main contributions of this work are as follows: (i) providing code for reproducing the reported results, (ii) subject dependent vs. subject-independent approaches, (iii) the ef-fect of different preprocessing steps (ICA, down-sampling and filtering), and (iv) achieving state-of-the-art performance on the 6 word classification task reporting a mean accuracy of 29.21% for all subjects on a publicly available dataset [8].

### 0.1. State-of-the-art Literature

Research studies in inner speech decoding use data of invasive (e.g., electrocorticography (ECoG) [9,10]) and non-invasive methods (e.g., magnetoencephalography (MEG) [11,12], functional magnetic resonance imaging (fMRI) [13], Functional near-infrared spectroscopy (FNIRS) [14,15]) with EEG being the most dominate modality used so far [16]. Martin et al. [10] attempted to detect single words from inner speech using ECoG recordings from inner and outer speech. This study included six word pairs and achieved a binary classification accuracy of 58% using a Support Vector Machine (SVM). ECoG is not scalable as it is invasive but it advances our understanding and limit of decoding inner speech research. Recent methods used a CNN with the “MEG-as-an-image” [12] and “EEG-as-raw-data” [7,17] inputs.

The focus of this paper is on inner speech decoding in terms of classification of the words and vowels. Classified words can be useful in many scenarios of human-computer communication, e.g., in smart homes or health-care devices, were the human wants to give simple commands by the brain signals in a natural way. For human-to-human communication, the ultimate goal of inner speech decoding (in terms of representation learning) is often to synthesize speech [18,19]. In this related area [18] uses a minimal invasive method called stereotactic EEG (sEEG) with one subject and 100 Dutch words, in an open-loop stage for training the decoding models and close-loop stage to evaluate in real time the imagined and whispered speech. The attempt, although not yet intelligible, provide a proof of concept for tackling close-loop synthesis of imagined speech in real time. [19] uses MEG data from 7 subjects, using as stimuli 5 phrases (1. Do you understand me, 2. That’s perfect, 3. How are you, 4. Good-bye, and 5. I need help.), and 2 words (yes/no). They follow a subject-dependent approach, where they train and tune a different model per subject. Using a bidirectional long short-term memory recurrent neural network, they achieve a correlation score of the reconstructed speech envelope of 0.41 for phrases and 0.77 for words.

[15] reported an average classification accuracy of 70.45 ± 19.19% for a binary word classification task using regularized linear discriminant analysis (RLDA) using FNIRS data. The EEGNet [20] is a CNN-based deep learning architecture for EEG signal analysis that includes a series of 2D convolutional layers, average pooling layers, and batch normalization layers with activations. Finally, there is a fully connected layer at the end of the network to classify the learned representations from preceding layers. The EEGNet serves as the backbone network in our model; however, the proposed model extends the EEGNet in similar manner to [7].

There are two main approaches, when it comes to brain data analysis: subject-dependent and subject-independent. In subject-dependent approach the analysis is taken for each subject individually and performance is reported per subject. Representative studies in the subject-dependent approach follows. [8] reported a mean recognition rate of 22.32% in classifying 5 vowels and 18.58% in classifying 6 words using a Random Forest (RF) algorithm, with a subject-dependent approach. Using data from 6 subjects, [21] reported an average accuracy of 50.1% ± 3.5 for the 3 word classification problem and 66.2% ± 4.8 for a binary classification problem (long vs. short words), following a subject-dependent approach using a multi-class Relevance Vector Machine (MRVM). In [12] MEG data from inner and outer speech and an average accuracy of 93% for the inner speech and 96% for the outer speech decoding of 5 phrases in a subject dependent approach using a CNN was reported. Recently, [22] reports an average accuracy of 29.7% for 4 words classification on a publicly available dataset of inner speech [23]. In subject-independent approach all subjects are taken into account and the performance is reported using data of all subjects. Therefore the generated decoding model can generalise in new subjects’ data.

The following studies uses a subject-independent approach. In [6], the authors reported an overall accuracy of 90% on the binary classification of vowels compared with consonants using Deep-Belief Networks (DBN) and the combination of all modalities (inner and outer speech), in a subject-independent approach. In [7], the authors used a CNN with transfer learning to analyse inner speech on the EEG dataset of [8]. In these experiments, the CNN was trained on the raw EEG data of all subjects but one. A subset of the remaining subject’s data was used to fine-tune the CNN and the rest of the data was used for testing the CNN model. They authors reported an overall accuracy of 35.68% (5-fold cross-validation) for the 5 vowel classification task.

## 1. Materials and Methods

### 1.1. Dataset and Experimental Protocol

The current work uses a publicly available EEG dataset as described in [8]. This dataset includes recordings from 15 subjects using their inner and outer speech to pronounce 5 vowels (*/a/, /e/, /i /, /o/, /u/*) and 6 words (*arriba/up, abajo/down, derecha/right, izquierda/left, adelante/forward, and atr as/backwards*). A total of 3316 and 4025 imagined speech sample EEG recordings for vowels and words, respectively, are available in the dataset. An EEG with 6 electrodes was used in these recordings.

Figure 1 shows the experimental design followed in [8]. The experimental protocol consisted of a ready interval that was presented for 2 sec, followed by the stimulus (vowel or word) presented for 2 sec. The subjects were asked to use their inner or outer speech during the imagine interval to pronounce the stimulus. Finally, a rest interval of 4 seconds was presented, indicating that the subjects could move or blink their eyes before proceeding with the next stimulus. It is important to note that for the purpose of our study, only the inner speech part of the experiment is used.

**Figure 1.**
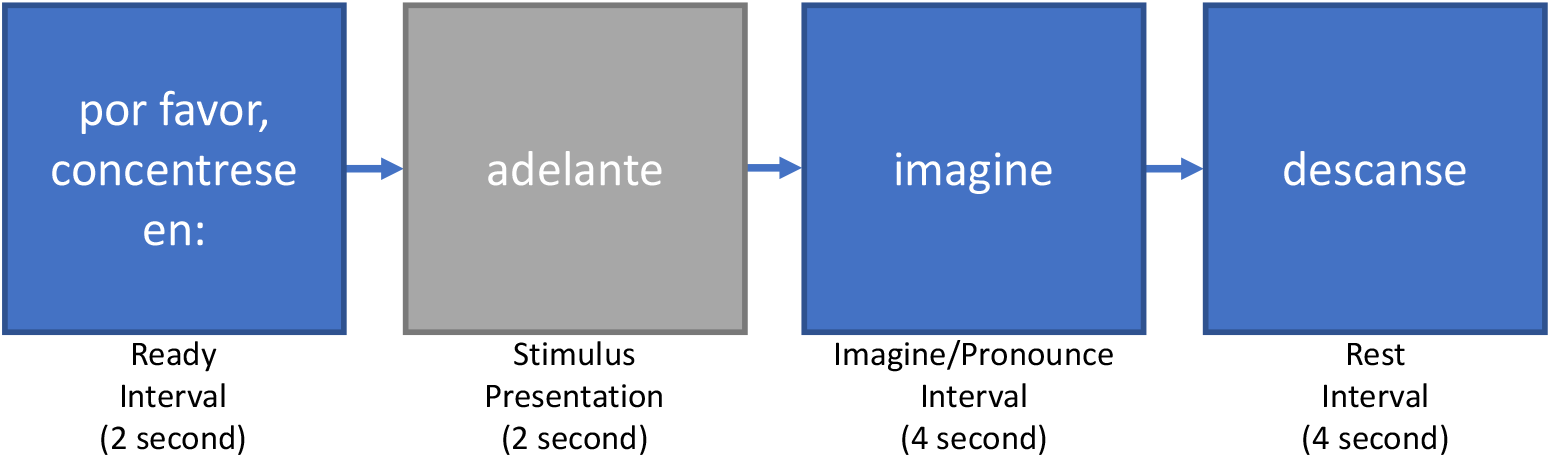
Experimental protocol used in [8]: Ready interval followed by a textual representation of the stimulus (vowel or word). The inner speech production took place during the stimulus interval for 4 seconds.

### 1.2. Methods

The proposed framework uses a deep CNN to extract representations from the input EEG signals. However, before applying the proposed CNN, the signals are preprocessed and then the CNN network is trained on the preprocessed signals.

Fig. 2 depicts the flow of the proposed work. Separate networks are trained for vowels and words following the architecture depicted in Fig. 2. The proposed network is inspired by Cooney et al. [7]; they performed filtering, downsampling, and artefact removal before applying the CNN. However, we have noticed that downsampling degrades the recognition performance, see Section 3. Therefore, we do not downsample the signals in our experiments. The downsampling block is represented by a cross in Fig. 2 to indicate that this task is not included in our proposed system in comparison with [7]. The current work reports results on 3 different experimental approaches using preprocessed data and raw data. The 3 different approaches are discussed in detail in Sections 2.1.2 - 2.1.1. More information about the preproccessing techniques can be found in Section 1.3.

**Figure 2.**
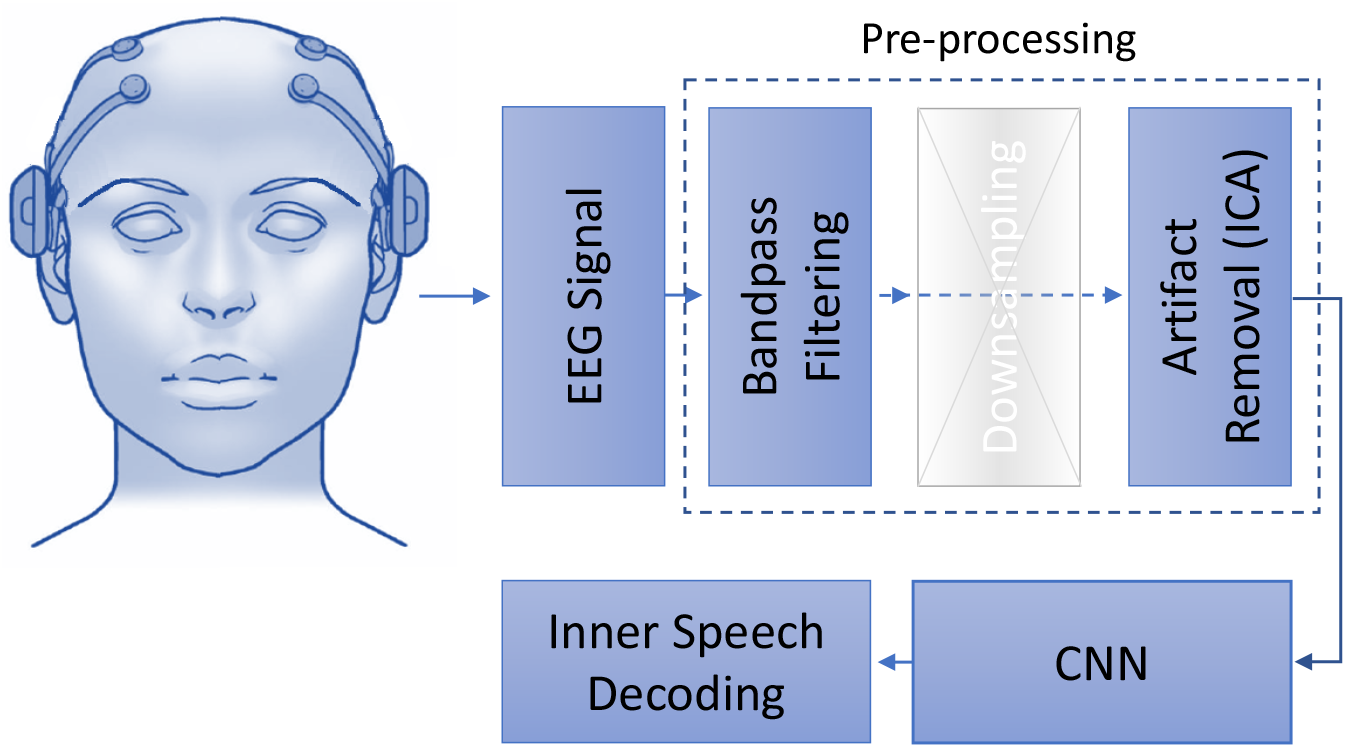
The figure illustrates the proposed workflow. The preprocessed EEG signals with or without downsampling are used to train a CNN model for inner speech decoding.

### 1.3. Preprocessing

In the current work we apply the following preprocessing steps:

#### Filtering

A frequency between 2 Hz and 40 Hz is used for filtering [8].

#### Down-sampling

The filtered data are down-sampled to 128 HZ. The original frequency of the data is 1024 Hz.

#### Artefact removal

Independent component analysis (ICA) is known as a blind-source separation technique. When recording a multi-channel signal, the advantages of employing ICA become most obvious. ICA facilitates the extraction of independent components from mixed signals by transforming a multivariate random signal. Here, ICA applied to identify components in EEG signal that include artefacts such as eye blinks or eye movements. These components then filtered out before the data is translated back from source space to sensor space. ICA effectively removes noise from the EEG data and is, therefore, an aid to classification. Given the small number of channels, we intact all the channels but used ICA [25] for artefact removal ^1^.

Figure A3 (see Appendix) depicts the preprocessed signal after applying ICA. This figure shows the vowel *a* for two subjects. From this figure, it can be noted that the subject’s model is not discriminative enough as overlapping is observed. The response from all electrodes’ behaviour for all vowels for Subject-02 can be seen in Figure A4 (see Appendix). From this figure, it can be seen that all electrodes are adding information as they all differ in their characteristics.

### 1.4. iSpeech-CNN Architecture

In this section, we introduce the proposed CNN-based iSpeech architecture. After extensive experiments on the existing CNN architecture for inner speech classification tasks, we determined that downsampling the signal has an effect on the accuracy of the classification and thus removed it from the proposed architecture. The iSpeech-CNN architecture for imagined vowel and word recognition is shown in Fig. 3. The same architecture is used in training for imagined vowels and words separately. The only difference is that the network for vowels has five classes, therefore, the softmax layer outputs five probability scores; one for each vowel. In the same manner, the network for words has six classes, therefore, the softmax layer outputs six probability scores; one for each word. Unlike [7], after extensive experimentation, we observed that the number of filters has an effect on the overall performance of the system; 40 filters are used in the first four layers of both networks. The next three layers have 100, 250, and 500 filters, respectively. However, the filter sizes are different. Filters of sizes (1 × 5), (6 × 1), (1 × 5), (1 × 3), (1 × 3), (1 × 3), and (1 × 3) are used in the first, second, third, fourth, fifth, sixth, and seventh layers, respectively.

**Figure 3.**
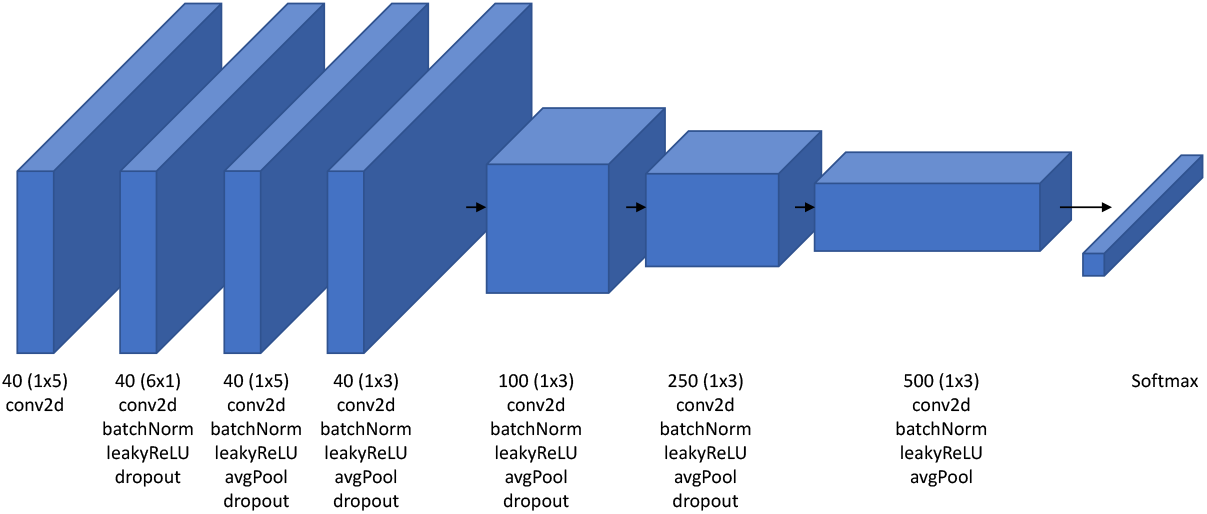
Proposed iSpeech-CNN architecture for imagined speech recognition based on the architecture described in [7]. This network is trained separately for vowels and words. Therefore, the difference lies in the last layer (softmax). The softmax layer for vowels have five outputs while for words has six outputs.

We used Adam optimizer with a dropout of 0.0002 for the vowel classification and 0.0001 for the word classification. As the network is very small, dropping out more features will adversely affect the performance. The initial learning rate was fixed to 0.001 with a *piecewise* learning rate scheduler. Our network was trained for 60 epochs, and the best validation loss was chosen for the final network. The regularization was also fixed to a value of 0.001. Our proposed iSpeech-CNN architecture follows the same structure as [7] but with a different numbers of filters and training parameters and preprocessing.

## 2. Experimental Approaches and Performance Measures

This section describes the experimental approaches that have been utilized for the analysis of EEG data and the performance measures that quantify the obtained analysis.

### 2.1. Experimental Approaches

Three experimental approaches were used for analysis, and they are discussed in detail in the following subsections.

#### 2.1.1. Subject-dependent/Within-subject Approach

Subject-dependent/within-subject classification is a baseline approach that is commonly used for the analysis of inner speech signals. In this approach, individual models are trained corresponding to each subject and for each subject, a separate model is created. The training, validation and testing sets will all have data from the same subject. This approach essentially measures how much an individual subject’s data changes (or varies) over time.

To divide the subject data into training, testing and validation datasets, a ratio of 80-10-10 is used. The training, validation and testing datasets contain all vowel/words category samples (five/six, respectively) in the mentioned ratio. To remove the bias towards the samples, five different trials are utilized. Furthermore, the mean accuracy and standard deviation are reported for all experimental approaches.

#### 2.1.2. Subject Independent: Leave-one-out Approach

The subject-dependent approach does not show generalization capability as it models a one subject at a time (Testing data contain only samples of the subject that is being modelled). The leave-one-out approach is an independent approach where data of each subject are tested using models that are trained using the data of all other subjects but one, i.e., *n* – 1 subjects out of total *n* will be used for the training model, and the rest will be used for testing. For example, Model-01 will be trained with data from subjects except Subject01, and will be tested with Subject01 (see Table [A3–A6]).

This approach helps to obtain a deeper analysis when there are fewer subjects or entities and shows how each individual subject affects the overall estimate of the rest of the subjects. Hence, this approach may provide more generalizable remarks than subject-specific models that depend on individual models.

#### 2.1.3. Mixed Approach

The mixed approach is a variation of subject-independent approach. While leave-one-out is truly independent, we can see mixed approach as less independent in nature as it includes data from all subjects in training, validation and testing. As, it contains the data of all subjects, we called it mixed approach. This approach differs from the within-subject and leave-one-out approaches, where *n* models, which corresponds to the total number of subjects in the data, are trained. In this approach, only one model will be trained for all subjects. Testing contains samples of all the subjects under all categories (vowels/words).

To run this experiment, 80% of the samples of all the subjects are included in the training set, 10% in the validation set and remaining in the test set. We also ensure the class balancing, i.e., each class will have approximately the same number of samples of all vowel/word categories. The same experiment is repeated for five random trials, and the mean accuracy along with the standard deviation is reported.

### 2.2. Performance Measures

The mean and standard deviation are used to report the performance of all the approaches. For the final results, the F-score are also given.

#### Mean

The mean is the average of a group of scores. The scores are totalled and then divided by the number of scores. The mean is sensitive to extreme scores when the population samples are small.

#### Standard deviation

In statistics, the standard deviation (SD) is a widely used measure of variability. It depicts the degree of deviation from the average (mean). A low SD implies that the data points are close to the mean, whereas a high SD suggests that the data span a wide range of values.

#### F-score

The F-score is a measure of a model’s accuracy that is calculated by combining the precision and recall of the model. It is calculated by the following formula:

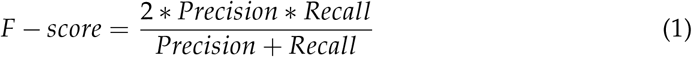

where precision is the percentage of true positive examples among the positive examples classified by the model, and recall is the fraction of examples classified as positive, among the total number of positive examples.

## 3. Results and Discussion: Vowels (5 Classes)

The results estimated with the subject-specific approach are discussed first as this approach is common in most of EEG related papers. All code, raw data, and preprocessed data are provided on Github ^2^. Related approaches are discussed in later subsections.

### 3.1. Subject-dependent/Within-subject Classification

In this section, we report the results when applying the subject-dependent approach. Figures A1, 5 and Table A5 show the results of our proposed iSpeech-CNN architecture. Table A1 and Table A2 show the results of the reference CNN architecture.

#### 3.1.1. Ablation study - influence of downsampling

Table A1 shows the results with raw and downsampled data when used within the referenced CNN architecture framework.

It is clearly observed from the Table A1 that downsampling signals results in loss of information. Figure 4 furthermore shows that there is a significant performance increase between 32 and 1024, however, some other differences (e.g., for 40 filters between 128 and 1024) are not significant. For clarity, the bars for standard error for each data point are added. The highest vowel recognition performance (35.20%) is observed at highest sampling rate (1024), i.e., without downsampling.

**Figure 4.**
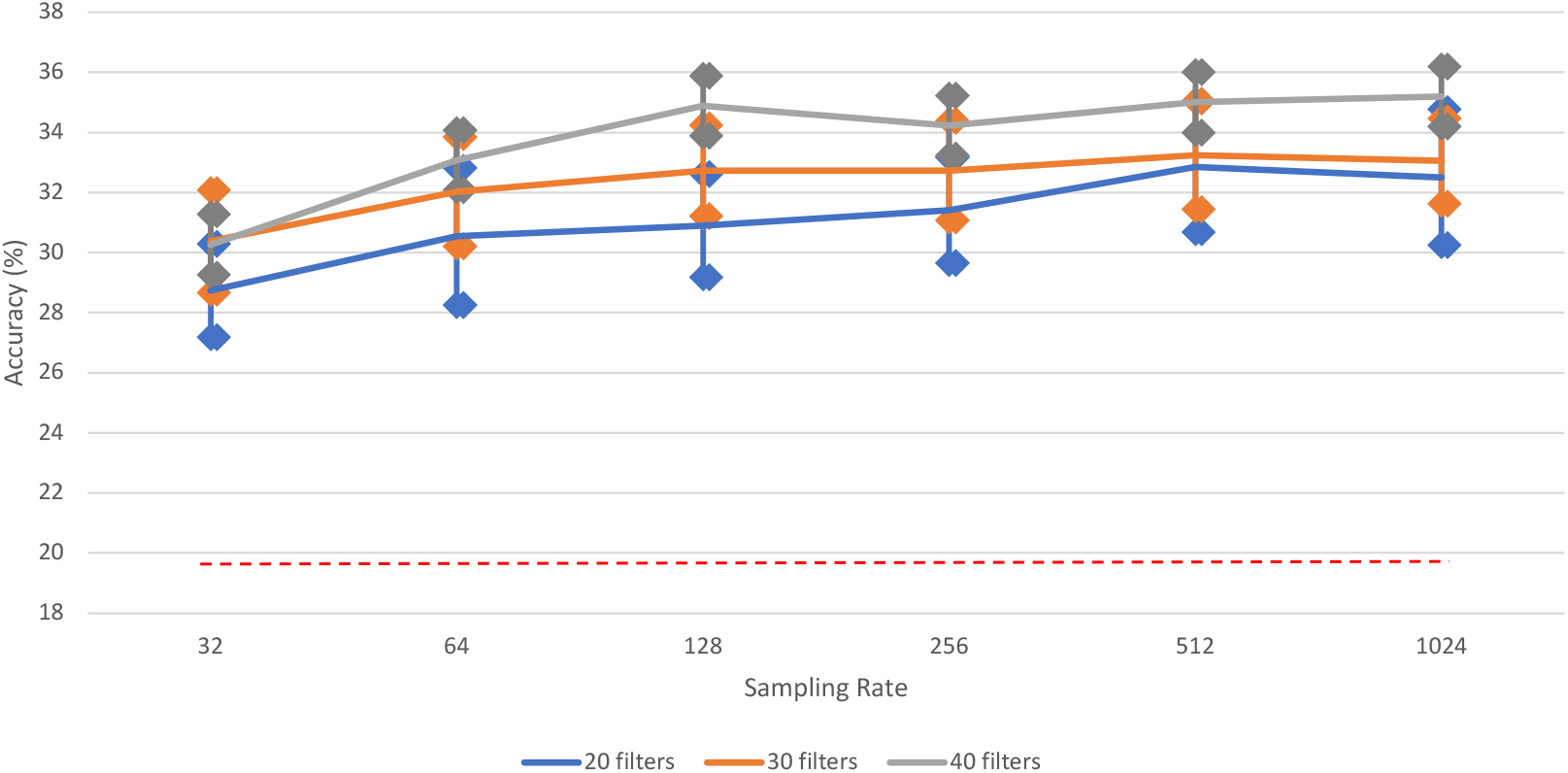
The impact of different sampling rates on vowel recognition performance of [iSpeech-CNN Architecture] with different filters in first three CNN layers. The bars indicate the standard error, sample size=5. Theoretical chance accuracy = 20% (red dotted line).

In other words, the chosen sampling rate was not sufficient enough to retain the original information. Therefore, further results will be reported for both raw data and downsampled data, in order to obtain a better insight into the preprocessing (i.e., filtering and ICA) stage.

#### 3.1.2. Ablation study - influence of preprocessing

Filtering and artefact removal plays an important role while analyzing the EEG signals. We applied the both bandpass filtering (see Section1), and picard (preconditioned ICA for real data) for artefact removal to obtain more informative signals. Table A2 shows the results of preprocessing when applied on the raw and downsampled data within the reference CNN architecture framework. The performance, i.e., the overall mean accuracy, decreased from 32.51% to 30.90%. The following points can be noted from Table A2: (1) Filtering and artefact removal highly influence the performance irrespective of raw and downsampled data. (2) The improved performance can also be observed with respect to each subject. A smaller standard deviation can also be seen. (3) The CNN framework generated higher performance than the handcrafted features, and the GRU (see Table 2). We also did experiments with LSTM classifier and noticed the random behavior (theoretical chance accuracies); no significant difference as compared to GRU. Therefore, iSpeech-CNN performs best among all classifiers.

**Table 1.**
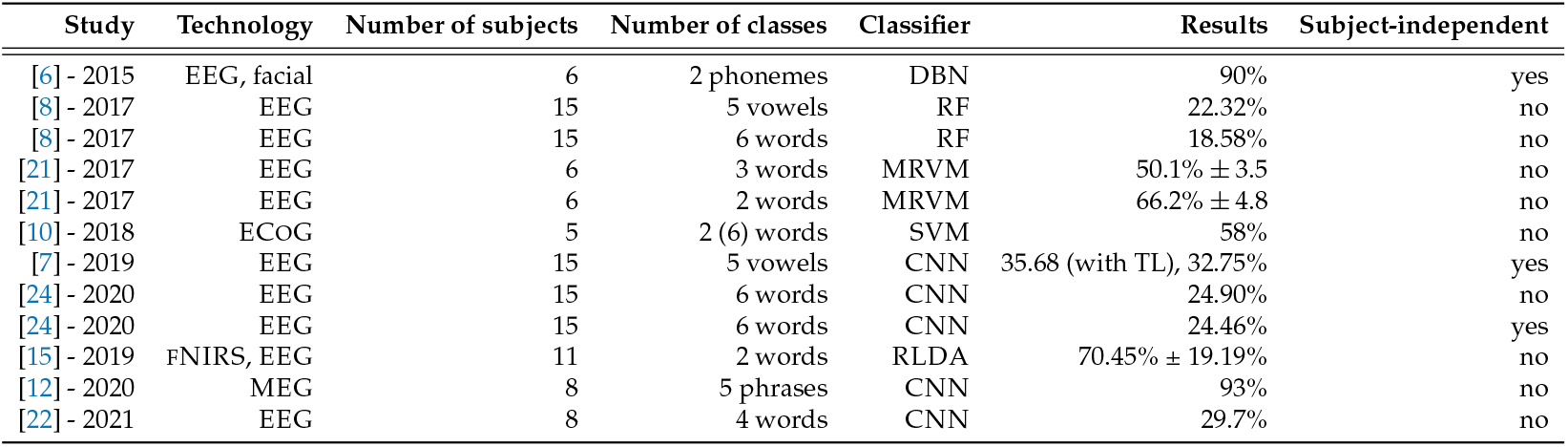
Overview of inner speech studies (2015-2021). TL: Transfer learning

| Study | Technology | Number of subjects | Number of classes | Classifier | Results | Subject-independent |
| --- | --- | --- | --- | --- | --- | --- |
| [6] - 2015 | EEG, facial | 6 | 2 phonemes | DBN | 90% | yes |
| [8] - 2017 | EEG | 15 | 5 vowels | RF | 22.32% | no |
| [8] - 2017 | EEG | 15 | 6 words | RF | 18.58% | no |
| [21] - 2017 | EEG | 6 | 3 words | MRVM | $50.1\% \pm 3.5$ | no |
| [21] - 2017 | EEG | 6 | 2 words | MRVM | $66.2\% \pm 4.8$ | no |
| [10] - 2018 | ECoG | 5 | 2 (6) words | SVM | 58% | no |
| [7] - 2019 | EEG | 15 | 5 vowels | CNN | 35.68 (with TL), 32.75% | yes |
| [24] - 2020 | EEG | 15 | 6 words | CNN | 24.90% | no |
| [24] - 2020 | EEG | 15 | 6 words | CNN | 24.46% | yes |
| [15] - 2019 | fNIRS, EEG | 11 | 2 words | RLDA | $70.45\% \pm 19.19\%$ | no |
| [12] - 2020 | MEG | 8 | 5 phrases | CNN | 93% | no |
| [22] - 2021 | EEG | 8 | 4 words | CNN | 29.7% | no |

**Table 2.** Average subject-dependent classification results on the [8] dataset

| Study | Classifier | Vowels | Words |
| --- | --- | --- | --- |
| [8] - 2017 | RF | 22.32%±1.81 | 18.58%±1.47 |
| [7,24] - 2019, 2020 | CNN | 32.75%±3.23 | 24.90%±0.93 |
| iSpeech-GRU | GRU | 19.28% ±2.15 | 17.28% ±1.45 |
| iSpeech-CNN<br>(proposed) | CNN | 35.20% ±3.99 | 29.21% ±3.12 |

#### 3.1.3. Ablation study - influence of architecture

Based on the CNN literature in the EEG paradigm [7] [26], adding more layers to the reference CNN architecture does not help to obtain improved performance. However, by changing the number of filters in the initial layers, some improvements can be observed. Based on the CNN literature for EEG signals, having a sufficient number of filters in the initial layers helps to obtain some improvement [7] [27]. Here, we choose three initial layers, unlike in natural images, in speech, initial layers are more specific to the task rather than the last few layers. The results with a changing number of filters in the initial layer within the iSpeech-CNN architecture are shown in Table A5. In the reference CNN architecture, this filter number was 20 for the initial three layers. However, we have changed this number to 40 (decided based on experimentation) in the iSpeech-CNN architecture. Table A5 clearly shows that changing the filter parameter gives higher performance than with the number of filters (compare to the reference architecture results in Tables A1 and A2 in the appendix). This improvement is observed with and without downsampled data and with respect to the subject (see Figure A1 and Figure 5). The standard deviation also decreases with these modifications (see Table A5).

**Figure 5.**
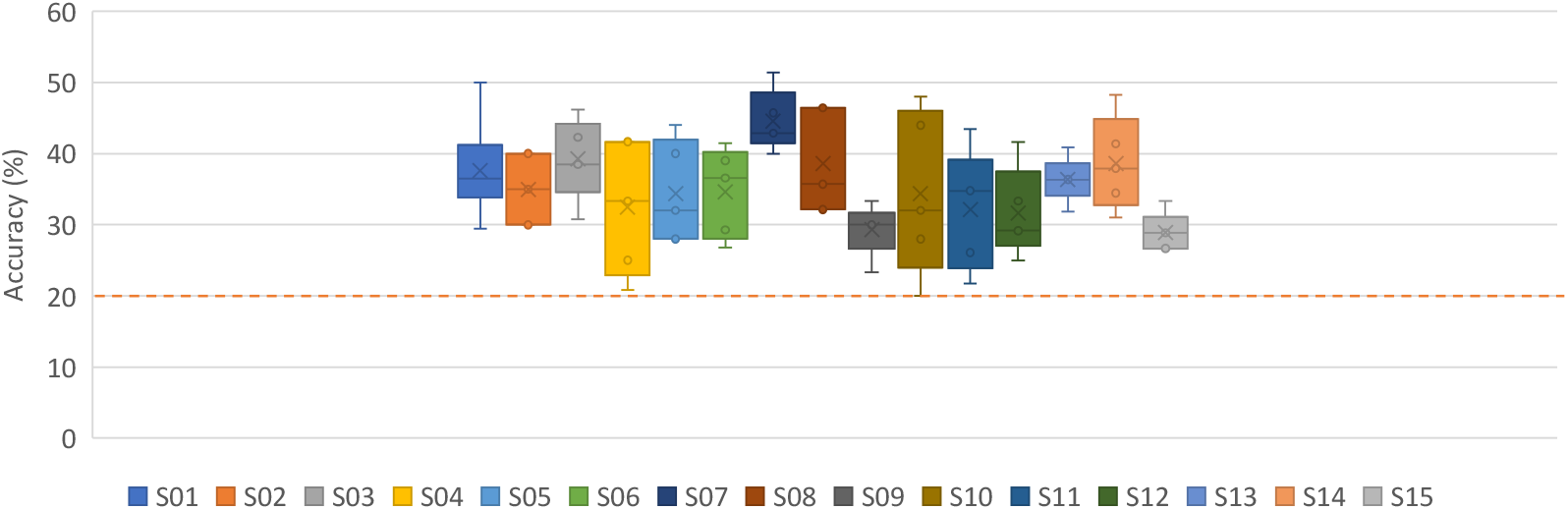
Subject dependent results for vowels without downsampling on preprocessed signals [iSpeech-CNN Architecture]. Theoretical chance accuracy = 20% (red dotted line).

### 3.2. Mixed approach results

This section discusses the results of mixed approach. In this approach, data from all subjects are included in training, validation and testing. Table A4 shows the results for the mixed approach with and without downsampling. These results were compiled with filtering and ICA in both reference and modified CNN architectures.

From these results, it is noted that the obtained accuracies are random in nature. The modified CNN architecture parameters, do not help to obtain any improvements, and show random accuracy behaviour. In other words, it is difficult to achieve generalized performance with EEG signals. Based on the EEG literature, it has also been justified that models trained on data from one subject cannot be generalize to other subjects even though have been recorded using the same setup conditions.

Determining the optimal frequency sub-bands corresponding to each subject could be one possible direction that may be successful in such a scenario. We intent to explore such a direction in our future work.

### 3.3. Subject-independent: Leave-one-out results

Having discussed the subject-specific and mixed results, in this section, the subject-independent results are discussed. The leave-one-out approach is a variation of the mixed approach, however, unlike the mixed approach, here the data of the testing subject are not included in the training. For example, in Figure 6, except *Subject*01, all other subjects were used in the training of *Model* – 01. Figure 6 and Table A6 show the results using the iSpeech-CNN architecture, while Table A3 shows the results using the reference architecture.

**Figure 6.**
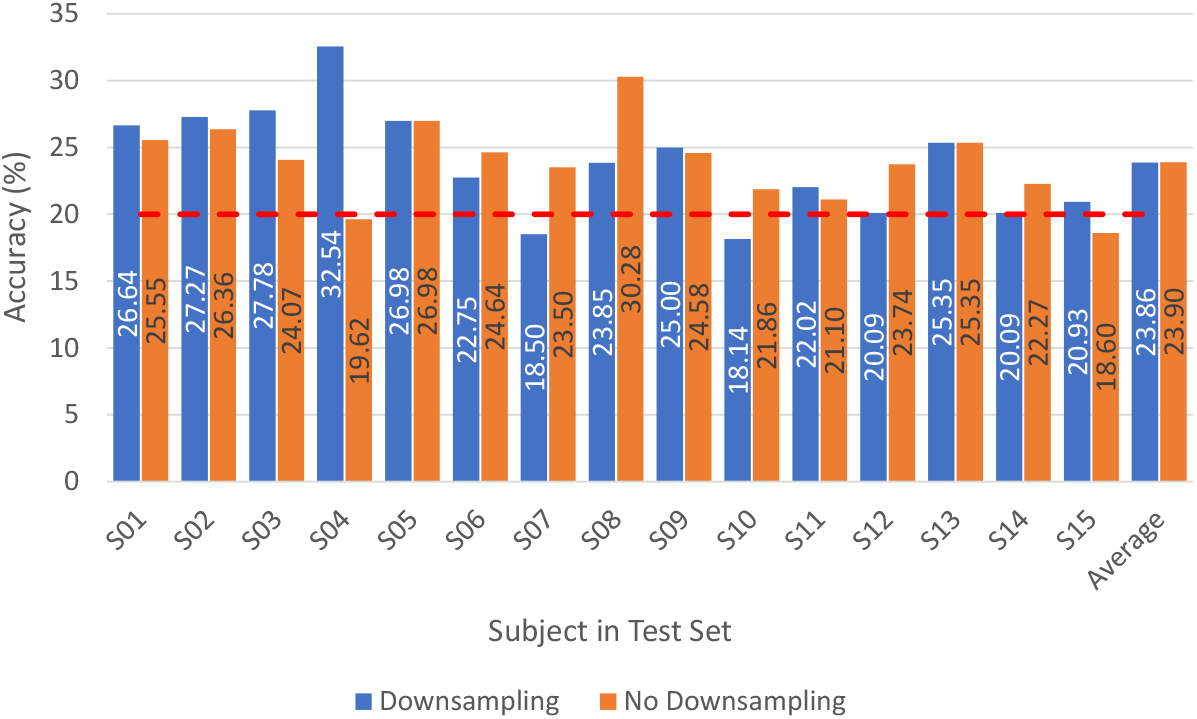
Leave-one-out results for vowels with and without downsampling on preprocessed signals [iSpeech-CNN Architecture]. Theoretical chance accuracy = 20% (red dotted line).

It can be noted that having fewer subjects in training (one less as compared to the mixed approach), shows slightly better behaviour than the mixed approach, where all subjects were included in the training. Moreover, changing the reference CNN parameters to our proposed iSpeech-CNN architecture also shows improved performance (see Figure 6 and Table A6).

The mixed and leave-one-out approaches both showed that generalizing the performance over all subjects is difficult in the EEG scenario. Hence, there is a need for the preprocessing stage, which can make the data more discriminative.

## 4. Results and Discussion: Words (6 Classes)

Having discussed all the approaches for the category of vowels, we noticed that only the subject-specific approach showed performance that was not random in nature and hence makes sense. Therefore, in this section, we only report results corresponding to the subject-specific approach for the word category.

This category contains six different classes (see Section 1.1). Table A8 and Figures A2 and 8 show the performance results for the classification of the 6 words, using the proposed iSpeech-CNN architecture. The performance results when using the reference architecture can be found in Appendix B. From these tables and figures, the same kind of behaviour as vowels is observed. The change in the number of filters in the initial layers affected the performance as shown in Table A8. The downsampling of data also affects the overall performance. Fig. 7 shows that the highest word recognition performance (29.12%) is observed at highest sampling rate (1024), i.e., without downsampling. For clarity, we added the bars for standard error for each data point. As opposed to vowel recognition, here is a steady increase of the performance when increasing the sampling rate (though again, not always significant among two neighboring values).

**Figure 7.**
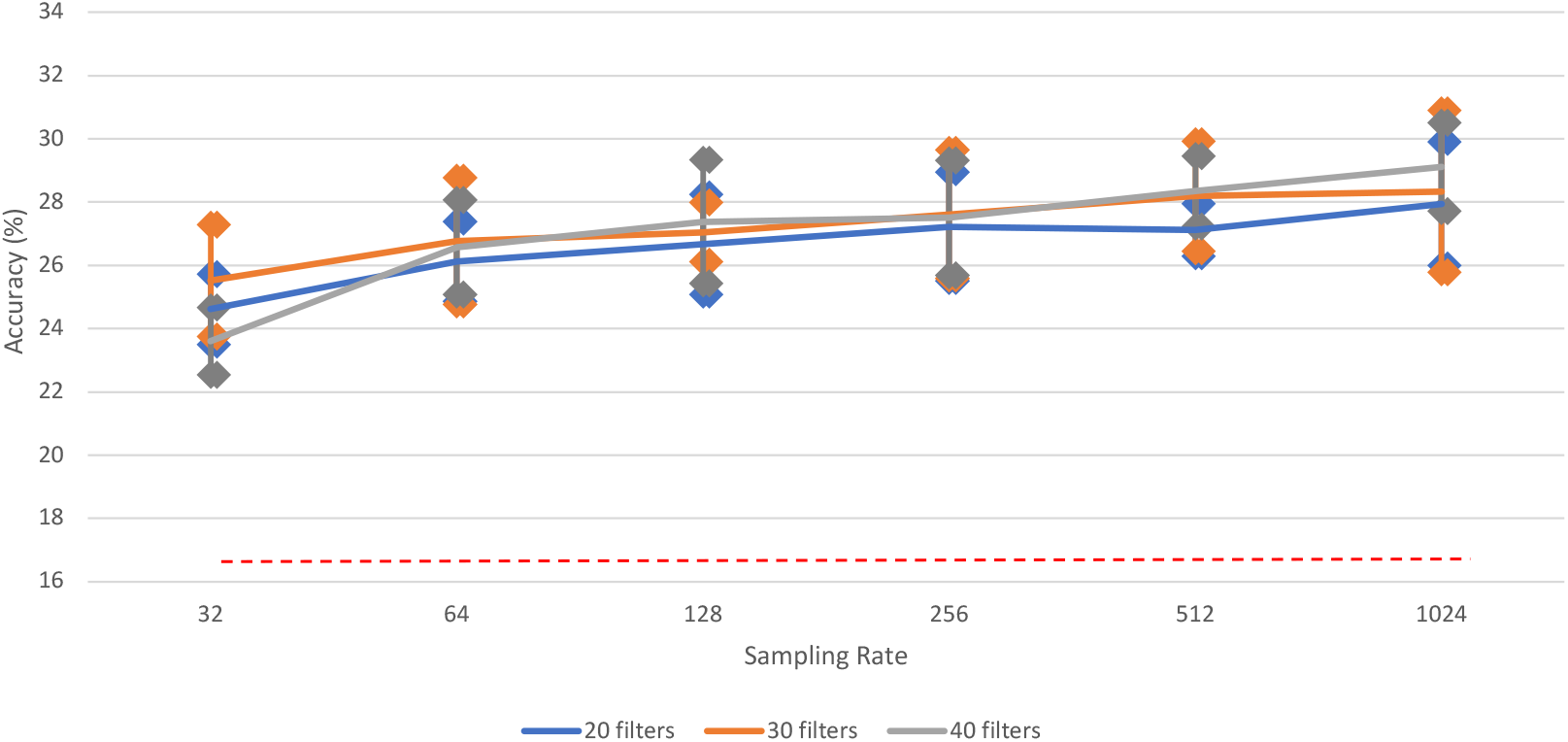
The impact of different sampling rates on word recognition performance of [iSpeech-CNN Architecture] with different filters in first three CNN layers. Performance increases with higher sampling rates. The bars indicate the standard error, sample size=5. Theoretical chance accuracy = 16.66% (red dotted line).

**Figure 8.**
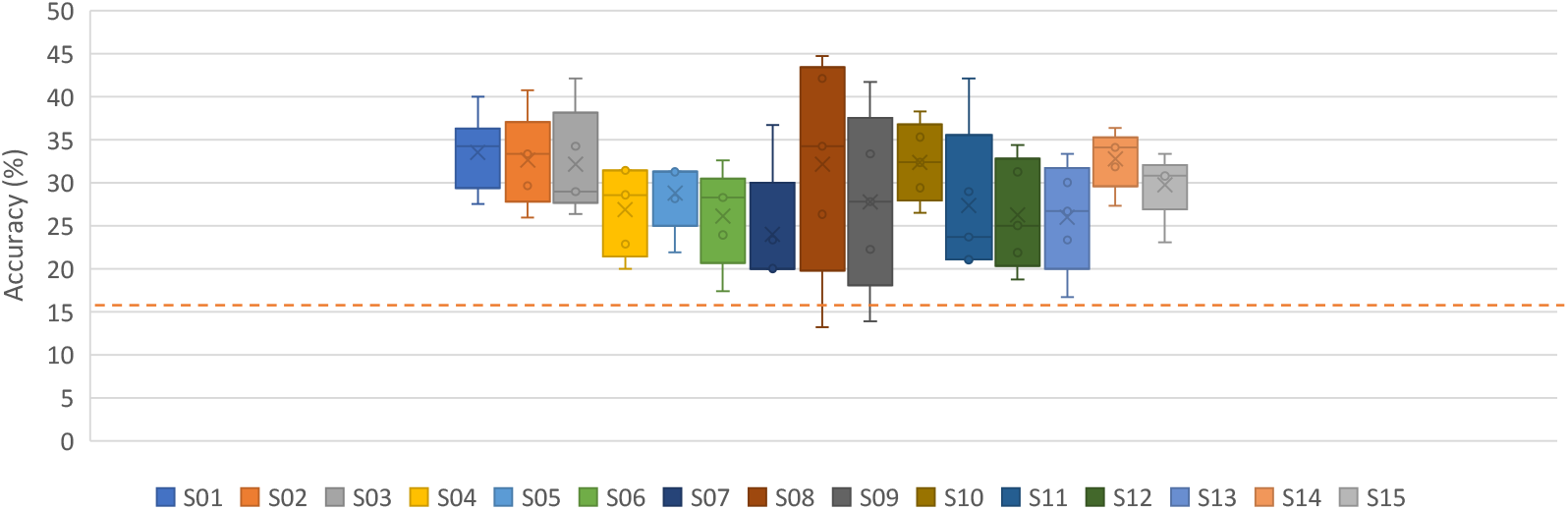
Subject-dependent results for words without downsampling on preprocessed signals [iSpeech-CNN Architecture]. Theoretical chance accuracy = 16.66% (red dotted line).

The iSpeech-CNN architecture shows better performance than handcrafted features such us real-time wavelet energy[8] and reference architecture (Appendix B).

Overall, we achieve a state-of-the-art performance of 29.21% when classifying the 6 words using our proposed iSpeech-CNN architecture and preprocessing methodology without downsampling.

The performance reported in this work is based on the CNN architecture of the reference network [7]. No other architecture was investigated. This is due to the reason that the goal of the proposed work is to reproduce the Cooney’s results and making the network and codes available to the research community.

## 5. Performance Comparison and related discussion

In this section, we compare our results on the vowels and words dataset with existing work and discuss related findings. Based on the reported performances in the Table 2, it is clearly noted that the CNN performs better than the handcrafted features for both datasets.

The precision, weighted F-score, and F-score for our proposed iSpeech-CNN in comparison with the reported results of Cooney et. al.[7] are shown in Table 3. From this table, we can note that our proposed system results in a higher precision however a lower F-score compared to the model in [7]. Hence, the reproducibility of the results reported in [7] is difficult.

**Table 3.** Precision and F-score (with respect to Tables A5–A8) for vowel and word classification [iSpeech-CNN Architecture].

| Vowel (iSpeech-CNN) |  |  |  |
| --- | --- | --- | --- |
|  | Precision | Weighted F score | F-score |
| No Downsampling | 34.85 | 41.12 | 28.45 |
| Downsampling | 34.62 | 38.99 | 30.02 |
| Cooney et al.[7] (Downsampling) | 33.00 | - | 33.17 |
| Words (iSpeech-CNN) |  |  |  |
|  | Precision | Weighted F-score | F-score |
| No Downsampling | 29.04 | 36.18 | 21.84 |
| Downsampling | 26.84 | 31.94 | 21.50 |

Our proposed CNN architecture and preprocessing methodology outperform the existing work in word and vowel category when following subject-dependent approach, as shown in Table 2. However it is worth to mention that for the vowel classification, unlike in [7], we don’t downsampling the data. Furthermore, [7] when using transfer learning approach for the vowel classification task, they report an overall accuracy of 35.68%, which is slightly higher than our reported accuracy in the subject-dependent approach.

Based on the 1-tail paired t-test results, we found that there is statistical significant difference between iSpeech-CNN and the reference paper [7] for word classification and for vowel classification, if we compare to the work without transfer learning (which is the fair comparison, as transfer learning adds a new dimension). We also found that there is no significant different between the best reported results with transfer learning [7,24] and iSpeech-CNN. Furthermore, when we run the 1-tail paired t-test results for iSpeech-CNN between downsampling and without downsampling, we found that these difference are significantly different for the words task (*p*=0.0005), but not statistically significant for the vowels task. We are following 1-tail paired t-test and used 10% of the overall samples, i.e., 332 for vowels and 403 for words.

Hence, it is observed that the correct selection of preprocessing methods and the number of filters in the CNN, greatly add to the performance. The elaborated results for each category and with each approach have been added to Appendices A and B.

## 6. Conclusions

This study explores the effectiveness of preprocessing steps and the correct selection of filters in the initial layers of the CNN in the context of both vowel and word classification. The classification results are reported on a publicly available inner speech dataset of 5 vowels and 6 words [8]. Based on the obtained accuracies, it is found that such a direction of exploration truly adds to the performance. We report state-of-the-art classification performance for vowels and words with mean accuracies of 35.20% and 29.21% respectively, without downsampling the original data. Mean accuracies of 34.88% and 27.38% have been reported for vowels and words, respectively, with downsampling. Furthermore, the proposed CNN code in this study is available to the public to ensure reproducibility of the research results and to promote open research. Our proposed iSpeech-CNN architecture and preprossessing methodology are the same for both datasets (vowels and words).

Evaluating our system in other publicly available datasets is part of our future work. Furthermore, we will address the issues related to the selection of the downsampling rate and the selection of the optimal frequency subband with respect to subjects.

## Appendix A. The results on vowels

**Table A1.** Subject-dependent results for the vowels using raw and downsampled data [Reference Architecture].

|  | Raw |  |  | Downsampled |  |  |
| --- | --- | --- | --- | --- | --- | --- |
|  | Train | Validation | Test | Train | Validation | Test |
| S01 | 91.07 | 36.00 | 35.29 | 79.07 | 39.20 | 26.47 |
| S02 | 92.44 | 29.00 | 17.00 | 72.00 | 24.00 | 16.00 |
| S03 | 83.65 | 34.00 | 27.69 | 86.94 | 26.00 | 19.23 |
| S04 | 89.94 | 38.00 | 33.33 | 89.33 | 30.00 | 31.67 |
| S05 | 86.59 | 16.00 | 21.60 | 72.82 | 19.00 | 17.60 |
| S06 | 91.73 | 27.00 | 25.85 | 80.00 | 29.00 | 27.80 |
| S07 | 95.17 | 31.00 | 26.86 | 85.66 | 33.00 | 28.00 |
| S08 | 88.12 | 29.00 | 21.43 | 83.06 | 24.00 | 21.43 |
| S09 | 83.89 | 35.20 | 30.00 | 82.38 | 30.40 | 26.00 |
| S10 | 88.35 | 29.00 | 20.80 | 63.88 | 26.00 | 19.20 |
| S11 | 80.91 | 25.00 | 22.61 | 86.06 | 23.00 | 16.52 |
| S12 | 88.57 | 29.00 | 26.67 | 96.23 | 37.00 | 30.83 |
| S13 | 80.00 | 38.00 | 29.09 | 93.94 | 36.00 | 31.82 |
| S14 | 79.66 | 27.20 | 22.76 | 73.49 | 27.20 | 23.45 |
| S15 | 83.20 | 23.00 | 20.00 | 91.73 | 28.00 | 20.89 |
| Average/Mean | 86.88 | 29.76 | 25.29 | 82.44 | 28.79 | 23.79 |
| Standard Deviation | 4.81 | 5.96 | 5.14 | 8.73 | 5.44 | 5.35 |

**Table A2.** Subject-dependent results for vowels with and without downsampling on preprocessed
data [Reference Architecture].

| Preprocessing (Filtering and Artefact Removal) |  |  |  |  |  |  |
| --- | --- | --- | --- | --- | --- | --- |
|  | No Downsampling |  |  | Downsampling |  |  |
|  | Train | Validation | Test | Train | Validation | Test |
| S01 | 96.65 | 38.40 | 38.82 | 91.44 | 43.20 | 38.82 |
| S02 | 92.78 | 40.00 | 25.00 | 99.33 | 32.00 | 27.00 |
| S03 | 99.88 | 38.00 | 34.62 | 99.41 | 36.00 | 32.31 |
| S04 | 99.15 | 43.00 | 32.50 | 97.45 | 42.00 | 30.83 |
| S05 | 89.29 | 37.00 | 26.40 | 93.41 | 34.00 | 35.20 |
| S06 | 95.47 | 38.00 | 37.56 | 98.80 | 38.00 | 32.20 |
| S07 | 98.62 | 36.00 | 27.43 | 81.52 | 28.00 | 28.00 |
| S08 | 97.06 | 45.00 | 38.57 | 98.59 | 39.00 | 32.14 |
| S09 | 97.41 | 36.00 | 35.33 | 97.41 | 32.00 | 30.00 |
| S10 | 96.24 | 35.00 | 37.60 | 98.47 | 35.00 | 28.80 |
| S11 | 88.69 | 30.00 | 31.30 | 91.66 | 35.00 | 33.91 |
| S12 | 86.86 | 35.00 | 30.83 | 91.31 | 28.00 | 24.17 |
| S13 | 99.54 | 40.00 | 33.64 | 95.89 | 46.00 | 31.82 |
| S14 | 87.43 | 39.20 | 35.86 | 84.11 | 40.00 | 33.79 |
| S15 | 91.87 | 33.00 | 22.22 | 99.73 | 30.00 | 24.44 |
| Average/Mean | 94.46 | 37.57 | 32.51 | 94.57 | 35.88 | 30.9 |
| Standard Deviation | 4.44 | 3.62 | 5.05 | 5.48 | 5.28 | 3.83 |

**Table A3.** Leave-one-out results for vowels with and without downsampling on preprocessed data [Reference Architecture]. Subject in test set.

| Preprocessing (Filtering and Artefact Removal) |  |  |  |  |
| --- | --- | --- | --- | --- |
|  | No Downsampling |  | Downsampling |  |
|  | Validation | Test | Validation | Test |
| S01 | 71.43 | 21.17 | 23.08 | 18.61 |
| S02 | 88.99 | 20.91 | 40.41 | 21.36 |
| S03 | 86.00 | 25.93 | 57.48 | 19.91 |
| S04 | 87.77 | 24.40 | 55.20 | 22.01 |
| S05 | 69.33 | 19.53 | 58.72 | 24.65 |
| S06 | 55.36 | 14.69 | 60.84 | 23.70 |
| S07 | 60.30 | 26.00 | 36.36 | 30.50 |
| S08 | 83.99 | 28.44 | 41.51 | 21.10 |
| S09 | 45.45 | 26.25 | 44.05 | 24.58 |
| S10 | 73.72 | 20.00 | 41.99 | 17.67 |
| S11 | 51.45 | 25.23 | 68.88 | 24.31 |
| S12 | 91.90 | 22.37 | 55.34 | 20.55 |
| S13 | 63.63 | 20.28 | 61.47 | 22.12 |
| S14 | 78.62 | 23.14 | 51.31 | 21.83 |
| S15 | 84.42 | 25.58 | 24.28 | 22.33 |
| Average/Mean | 72.82 | 22.93 | 48.06 | 22.35 |
| Standard Deviation | 14.84 | 3.55 | 13.10 | 2.95 |

**Table A4.** Mixed-approach results for vowels with and without downsampling on preprocessed data (Filtering and Artefact Removal) [Reference Architecture; iSpeech-CNN Architecture].

| With preprocessing; Reference architecture parameters |  |  |  |  |  |  |
| --- | --- | --- | --- | --- | --- | --- |
|  | No Downsampling |  |  | Downsampling |  |  |
|  | Train | Validation | Test | Train | Validation | Test |
| **Trial 1** | 72.45 | 20.95 | 22.27 | 62.61 | 20.32 | 17.63 |
| **Trial 2** | 76.73 | 22.22 | 19.03 | 58.75 | 20.63 | 24.36 |
| **Trial 3** | 64.82 | 22.54 | 19.26 | 50.58 | 20.32 | 20.65 |
| **Trial 4** | 67.55 | 18.10 | 19.95 | 57.04 | 18.41 | 20.19 |
| **Trial 5** | 60.78 | 20.32 | 22.04 | 47.78 | 22.86 | 23.90 |
| **Mean/Average** | **68.47** | **20.83** | **20.51** | **55.35** | **20.51** | **21.35** |
| **Standard Deviation** | **5.61** | **1.59** | **1.38** | **5.43** | **1.42** | **2.50** |
| With preprocessing; iSpeech-CNN Architecture parameters |  |  |  |  |  |  |
|  | no Downsampling |  |  | Downsampling |  |  |
|  | Train | Validation | Test | Train | Validation | Test |
| **Trial 1** | 57.98 | 20.63 | 20.42 | 55.10 | 21.90 | 18.33 |
| **Trial 2** | 68.79 | 21.27 | 19.72 | 46.46 | 23.17 | 22.04 |
| **Trial 3** | 54.63 | 21.59 | 21.11 | 44.44 | 18.73 | 22.27 |
| **Trial 4** | 37.70 | 17.46 | 20.42 | 24.36 | 21.27 | 21.35 |
| **Trial 5** | 86.15 | 24.13 | 20.19 | 57.20 | 22.86 | 20.42 |
| **Average/Mean** | **61.05** | **21.02** | **20.37** | **45.51** | **21.59** | **20.88** |
| **Standard Deviation** | **16.04** | **2.14** | **0.45** | **11.64** | **1.58** | **1.43** |

**Table A5.** Subject dependent results for vowels with and without downsampling on preprocessed signals [iSpeech-CNN Architecture].

| Preprocessing (Filtering and Artefact Removal) |  |  |  |  |  |  |
| --- | --- | --- | --- | --- | --- | --- |
|  | No Downsampling |  |  | Downsampling |  |  |
|  | Train | Validation | Test | Train | Validation | Test |
| **S01** | 86.33 | 48.80 | 37.65 | 81.86 | 33.60 | 37.65 |
| **S02** | 98.00 | 38.00 | 35.00 | 84.33 | 31.00 | 30.00 |
| **S03** | 85.76 | 38.00 | 39.23 | 98.00 | 37.00 | 36.15 |
| **S04** | 97.09 | 46.00 | 32.50 | 97.45 | 41.00 | 35.83 |
| **S05** | 91.41 | 44.00 | 34.40 | 94.59 | 38.00 | 36.00 |
| **S06** | 89.87 | 41.00 | 34.63 | 95.33 | 38.00 | 31.71 |
| **S07** | 98.34 | 39.00 | 44.57 | 98.21 | 39.00 | 41.14 |
| **S08** | 80.24 | 41.00 | 38.57 | 96.82 | 38.00 | 38.57 |
| **S09** | 96.65 | 32.80 | 29.33 | 92.22 | 35.20 | 30.00 |
| **S10** | 97.41 | 44.00 | 34.40 | 87.53 | 37.00 | 40.80 |
| **S11** | 95.43 | 41.00 | 32.17 | 98.06 | 31.00 | 30.43 |
| **S12** | 91.54 | 40.00 | 31.67 | 83.43 | 34.00 | 35.00 |
| **S13** | 82.74 | 46.00 | 36.36 | 85.60 | 46.00 | 37.27 |
| **S14** | 80.23 | 43.20 | 38.62 | 78.06 | 36.80 | 33.79 |
| **S15** | 89.60 | 31.00 | 28.89 | 90.80 | 38.00 | 28.89 |
| **Average/Mean** | **90.71** | **40.92** | **35.20** | **90.82** | **36.91** | **34.88** |
| **Standard Deviation** | **6.24** | **4.65** | **3.99** | **6.60** | **3.66** | **3.83** |

**Table A6.** Leave-one-out results for vowels with and without downsampling on preprocessed signals [iSpeech-CNN Architecture].

| **With Preprocessing (Filtering and Artefact Removal)** |  |  |  |  |
| --- | --- | --- | --- | --- |
|  | **No Downsampling** |  | **Downsampling** |  |
|  | **Validation** | **Test** | **Validation** | **Test** |
| **S01** | 46.65 | 26.64 | 38.23 | 25.55 |
| **S02** | 83.11 | 27.27 | 42.02 | 26.36 |
| **S03** | 74.32 | 27.78 | 43.16 | 24.07 |
| **S04** | 22.27 | 32.54 | 56.16 | 19.62 |
| **S05** | 64.79 | 26.98 | 56.24 | 26.98 |
| **S06** | 77.13 | 22.75 | 47.73 | 24.64 |
| **S07** | 47.21 | 18.50 | 40.76 | 23.50 |
| **S08** | 39.86 | 23.85 | 50.16 | 30.28 |
| **S09** | 82.38 | 25.00 | 46.36 | 24.58 |
| **S10** | 45.82 | 18.14 | 48.02 | 21.86 |
| **S11** | 52.42 | 22.02 | 54.23 | 21.10 |
| **S12** | 34.26 | 20.09 | 67.03 | 23.74 |
| **S13** | 46.05 | 25.35 | 50.08 | 25.35 |
| **S14** | 46.94 | 20.09 | 41.85 | 22.27 |
| **S15** | 51.69 | 20.93 | 46.31 | 18.60 |
| **Average/Mean** | **54.33** | **23.86** | **48.56** | **23.90** |
| **Standard Deviation** | **18.13** | **4.02** | **7.51** | **2.97** |

**Figure A1.**
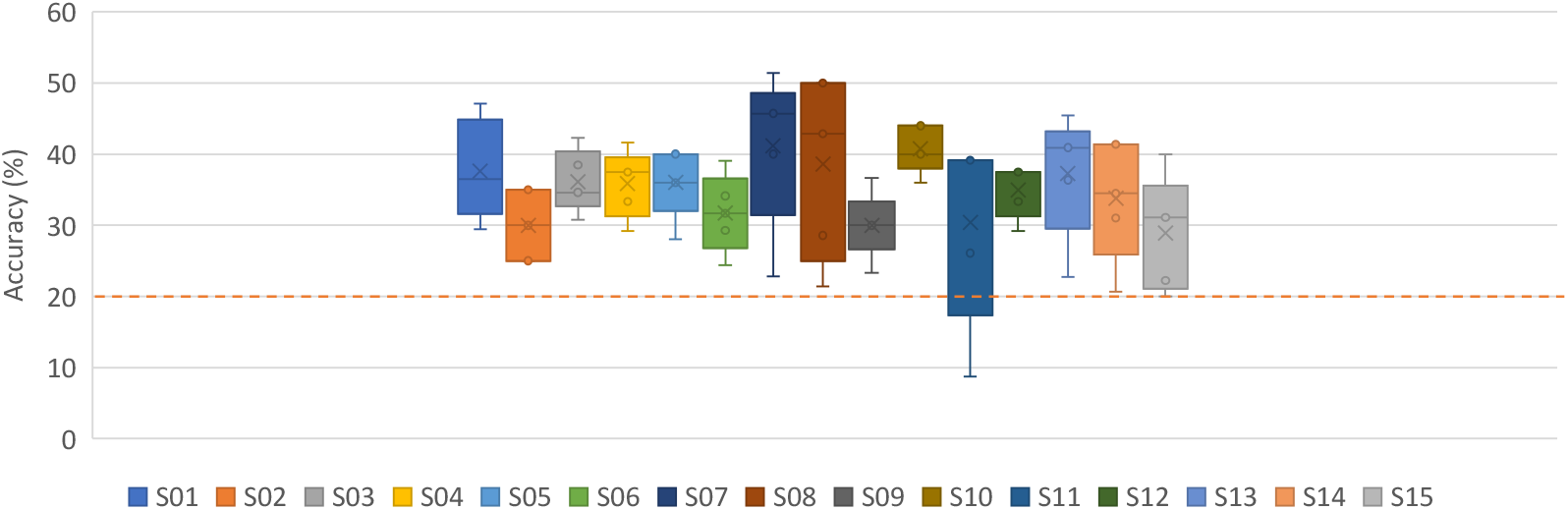
Subject dependent results for vowels with downsampling on preprocessed signals [iSpeech-CNN Architecture]. Chance accuracy 20%.

## Appendix B. The results on words

**Table A7.** Subject-dependent results for words with and without downsampling on preprocessed data [Reference Architecture].

| Preprocessing (Filtering and Artefact Removal) |  |  |  |  |  |  |
| --- | --- | --- | --- | --- | --- | --- |
|  | No Downsampling |  |  | With Downsampling |  |  |
|  | Train | Validation | Test | Train | Validation | Test |
| S01 | 90.60 | 31.33 | 30.00 | 83.16 | 32.67 | 28.50 |
| S02 | 96.57 | 30.83 | 23.70 | 79.62 | 30.00 | 22.22 |
| S03 | 90.19 | 32.00 | 31.58 | 67.33 | 30.00 | 29.47 |
| S04 | 92.75 | 36.67 | 25.14 | 75.49 | 33.33 | 25.71 |
| S05 | 87.53 | 41.67 | 28.13 | 69.89 | 30.83 | 31.25 |
| S06 | 83.12 | 39.17 | 25.65 | 71.72 | 29.17 | 23.04 |
| S07 | 92.22 | 26.67 | 25.33 | 79.39 | 35.00 | 21.33 |
| S08 | 85.37 | 38.33 | 30.53 | 86.48 | 34.17 | 23.16 |
| S09 | 92.86 | 32.50 | 25.56 | 82.57 | 30.00 | 28.89 |
| S10 | 89.05 | 31.67 | 29.41 | 92.00 | 30.00 | 30.59 |
| S11 | 93.14 | 31.33 | 27.37 | 95.24 | 32.00 | 22.11 |
| S12 | 88.00 | 33.33 | 25.62 | 82.00 | 34.67 | 32.50 |
| S13 | 96.76 | 28.33 | 33.33 | 86.10 | 39.17 | 25.33 |
| S14 | 86.10 | 35.33 | 28.64 | 68.95 | 36.00 | 28.18 |
| S15 | 96.98 | 27.50 | 29.23 | 91.77 | 28.33 | 27.69 |
| Average/Mean | 90.75 | 33.11 | 27.95 | 80.78 | 32.36 | 26.66 |
| Standard Deviation | 4.27 | 4.36 | 2.76 | 8.48 | 2.91 | 3.53 |

**Table A8.** Subject-dependent results for words with and without downsampling on preprocessed signals [iSpeech-CNN Architecture].

| Preprocessing (Filtering and Artefact Removal) |  |  |  |  |  |  |
| --- | --- | --- | --- | --- | --- | --- |
|  | No Downsampling |  |  | Downsampling |  |  |
|  | Train | Validation | Test | Train | Validation | Test |
| S01 | 97.95 | 34.67 | 33.50 | 77.35 | 30.00 | 25.50 |
| S02 | 92.86 | 34.17 | 32.59 | 89.05 | 28.33 | 23.70 |
| S03 | 93.05 | 32.00 | 32.11 | 76.67 | 28.67 | 34.74 |
| S04 | 94.51 | 35.00 | 26.86 | 95.59 | 29.17 | 31.43 |
| S05 | 95.38 | 34.17 | 28.75 | 82.47 | 26.67 | 31.25 |
| S06 | 96.99 | 38.33 | 26.09 | 67.20 | 30.83 | 26.09 |
| S07 | 87.47 | 33.33 | 24.00 | 80.71 | 30.00 | 16.00 |
| S08 | 98.33 | 35.00 | 32.11 | 83.70 | 30.00 | 28.95 |
| S09 | 98.86 | 37.50 | 27.78 | 70.00 | 27.50 | 28.89 |
| S10 | 92.38 | 32.50 | 32.35 | 93.52 | 34.17 | 27.06 |
| S11 | 97.43 | 36.67 | 27.37 | 87.90 | 35.33 | 26.32 |
| S12 | 92.86 | 37.33 | 26.25 | 69.05 | 32.00 | 25.62 |
| S13 | 84.57 | 31.67 | 26.00 | 68.29 | 31.67 | 33.33 |
| S14 | 96.38 | 37.33 | 32.73 | 66.38 | 30.67 | 27.27 |
| S15 | 94.69 | 33.33 | 29.74 | 86.67 | 31.67 | 24.62 |
| Average/Mean | 94.25 | 34.87 | 29.21 | 79.64 | 30.45 | 27.38 |
| Standard Deviation | 4.00 | 2.14 | 3.12 | 9.51 | 2.25 | 4.37 |

**Figure A2.**
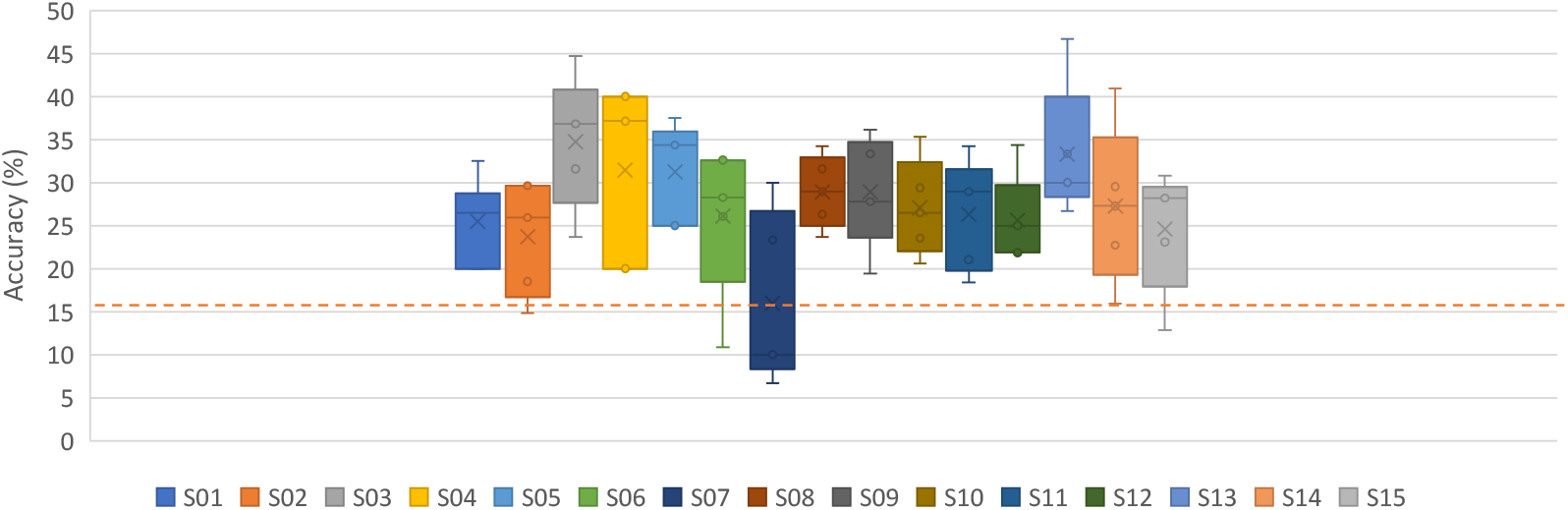
Subject-dependent results for words with downsampling on preprocessed signals [iSpeech-CNN Architecture]. Chance accuracy 16.66%.

## Appendix C. Dataset samples

**Figure A3.**
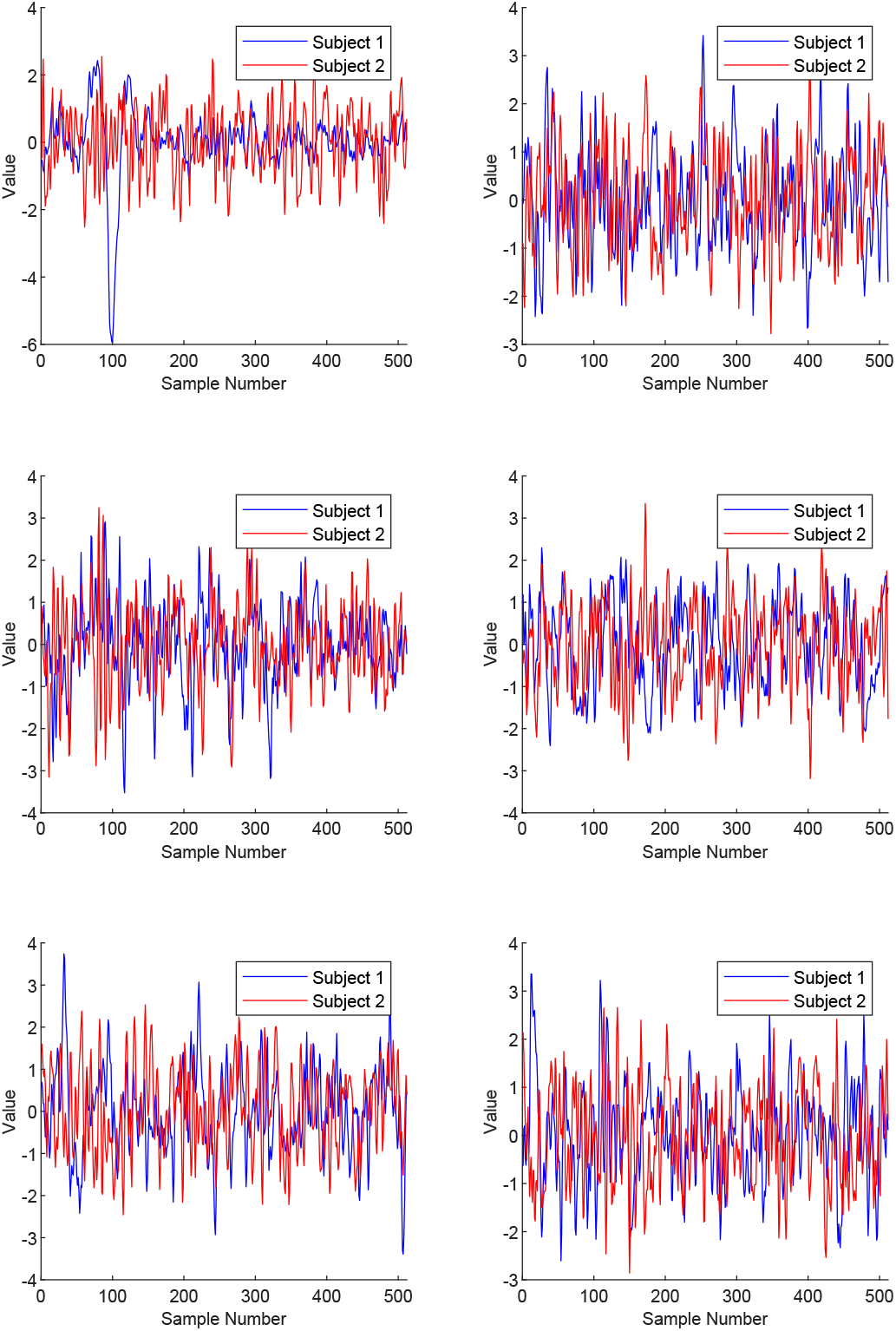
Example of preprocessed signals for all electrodes (after ICA) for the vowel /a/ for Subject01 and Subject02.

**Figure A4.**
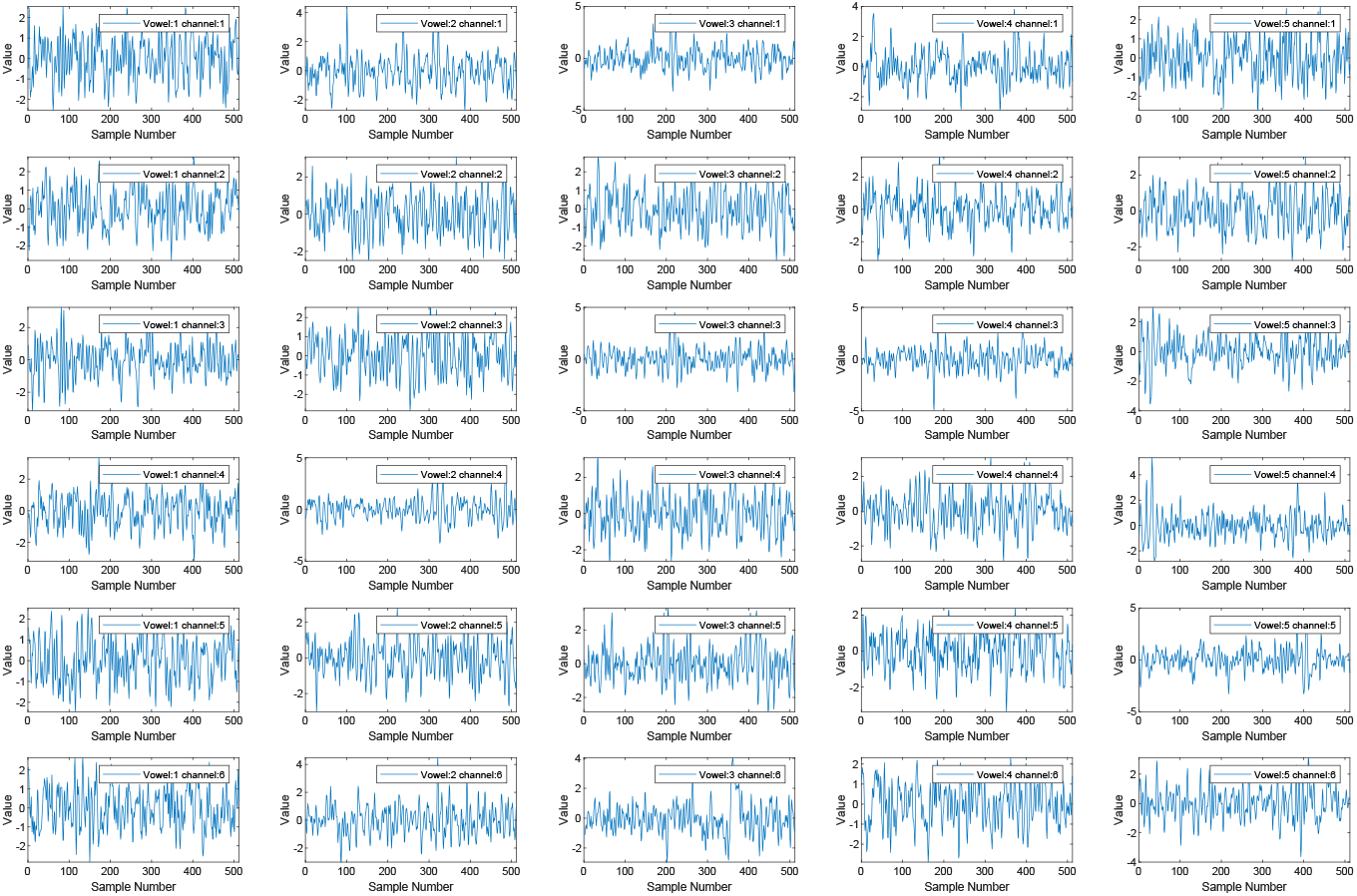
Example of preprocessed signals (after ICA) for all vowels and all electrodes for Subject02.

## Footnotes

1 https://github.com/pierreablin/picard/blob/master/matlab_octave/picard.m

2 https://github.com/LTU-Machine-Learning/Rethinking-Methods-Inner-Speech

